# Bringing Plant Immunity to Light: A Genetically Encoded, Bioluminescent Reporter of Pattern Triggered Immunity in *Nicotiana benthamiana*

**DOI:** 10.1101/2022.07.28.501876

**Authors:** Anthony G. K. Garcia, Adam D. Steinbrenner

## Abstract

Plants rely on innate immune systems to defend against a wide variety of biotic attackers. Key components of innate immunity include cell-surface pattern recognition receptors (PRRs), which recognize pest/pathogen-associated molecular patterns (PAMPs). Unlike other classes of receptors which often have visible cell death immune outputs upon activation, PRRs generally lack rapid methods for assessing function. Here, we describe a genetically encoded bioluminescent reporter of immune activation by heterologously-expressed PRRs in the model organism *Nicotiana benthamiana.* We characterized *N. benthamiana* transcriptome changes in response to *Agrobacterium tumefaciens (Agrobacterium)* and subsequent PAMP treatment to identify PTI-associated marker genes, which were then used to generate promoter-luciferase fusion fungal bioluminescence pathway (FBP) constructs. A reporter construct termed *pFBP_2xNbLYS1::LUZ* allows for robust detection of PTI activation by heterologously expressed PRRs. Consistent with known PTI signaling pathways, activation by receptor-like protein (RLP) PRRs is dependent on the known adaptor of RLP PRRs, SOBIR1. This system minimizes the amount of labor, reagents, and time needed to assay function of PRRs and displays robust sensitivity at biologically relevant PAMP concentrations, making it ideal for high throughput screens. The tools described in this paper will be powerful for studying PRR function and investigations to characterize the structure-function of plant cell surface receptors.

## Introduction

Plants perceive pests and pathogens through cell surface-localized immune receptors, termed pattern recognition receptors (PRRs). Canonically, these transmembrane proteins activate pattern triggered immunity (PTI) in response to conserved pathogen associated molecular patterns (PAMPs) (Boutrot & Zipfel, 2017). PTI consists of a suite of defense signaling and outputs including reactive oxygen species (ROS) production, ethylene production, peroxidase upregulation, callose deposition, stomatal modifications, calcium oscillations, and phytohormone production (Aldon et al., 2018; Berens et al., 2017; Broekgaarden et al., 2015; Melotto et al., 2017; Mott et al., 2018; Qi et al., 2017; Toyota et al., 2018; Y. Wang et al., 2021). These outputs aid in transcriptional reprogramming to improve plant resistance against attackers (Denoux et al., 2008; Navarro et al., 2004). Understanding immune activation by PRRs is critical for developing novel strategies to improve plant resistance against pests and pathogens.

The model organism *Nicotiana benthamiana* represents a significant resource in the field of plant immunity, in part because of robust immune phenotypes conferred by transiently expressed intracellular plant immune receptors (Goodin et al., 2008; Buscaill et al., 2021). *Agrobacterium-mediated* transient transformation of N. *benthamiana* allows for rapid expression of proteins, which is particularly applicable for mutant screening and structure-function analysis. For example, screening cell death as a visual reporter triggered by NLR activation has allowed investigations of structural features of nucleotide-binding leucine-rich repeat (NLR) proteins (Segretin et al., 2014; Steinbrenner et al., 2015; Adachi et al., 2019).

Transient transformation of *N. benthamiana* similarly serves as a powerful tool for studying PRRs, but there is currently a lack of robust visual reporters of PRR function in *N. benthamiana* analogous to cell death. Several cell surface immune receptors activate cell death phenotypes in other species, including leucine-rich repeat receptor-like proteins (LRR-RLPs) such as *Arabidopsis thaliana* RLP42 and *Solanum lycopersicum* Ve1, but these LRR-RLPs do not necessarily activate cell death upon heterologous expression in *N. benthamiana* or can require strong repeated elicitation (de Jonge et al., 2012; Z. Zhang et al., 2013; L. Zhang et al., 2014). Because heterologously expressed PRRs do not activate visual markers of PTI in *N. benthamiana,* immune responses mediated by transiently expressed PRRs are instead detected using early markers of PTI defense activation, including PAMP-induced ROS, ethylene, or peroxidase production (Mott et al., 2018; Steinbrenner et al., 2020). However, these assays are laborious or are hampered by the presence of *Agrobacterium* as a background source of PTI activation.

Reporters utilizing luminescence, fluorescence, or pigmentation have been adapted to study a variety of plant signaling processes (DeBlasio et al., 2010; Furuhata et al., 2020; He et al., 2020). However, no transiently expressed reporters of immune activation in intact *N. benthamiana* leaves have been described. A high sensitivity luciferase-based system for measuring pattern triggered immunity in protoplasts of *N. benthamiana* was previously reported (Nguyen et al., 2010), but was not tested for heterologously expressed PRRs and required external addition of luciferin. A different system using a bioluminescent strain of *Agrobacterium* expressing the bacterial *lux* operon allows for monitoring of *Agrobacterium* during transient transformation and quantification of effector triggered immunity (ETI), but has not yet been applied to PTI (Jutras et al., 2021). *N. benthamiana* lines stably expressing the fluorescent Ca2+ indicator GCaMP3 allow for detection of signaling in response to various biotic and abiotic stresses, including signaling activated by transiently expressed receptor-like kinases (DeFalco et al., 2017). However, stable expression of GCaMP3 limits the ability to quickly test different *N. benthamiana* genotypes and mutants lacking components of the signaling pathway. Finally, fluorescent proteins and pigments are simple to measure, but lack the same low background and high sensitivity of luciferase-based assays (Haugwitz et al., 2008; Thorne et al., 2010), which limits the ability to detect the range of responses that may occur in response to an immune elicitor.

To develop a generic PTI reporter, we performed transcriptomic analysis of *N. benthamiana* upon activation of a heterologously expressed PRR and adapted endogenous markers into a luciferase-based system that retains sensitivity but eliminates the need to introduce exogenous substrate. By encoding a metabolic pathway that allows for endogenous production of fungal luciferin alongside the fungal luciferase *(LUZ)* enzyme, the fungal bioluminescence pathway (FBP) system circumvents requirements for external addition of substrate while still remaining sensitive to subtle changes in gene expression (Khakhar et al., 2020; Mitiouchkina et al., 2020). Importantly, the features of the FBP system were well-suited for a reporter system that meets several criteria to be useful for studying plant cell surface immune receptors in a heterologous system: 1) highly sensitive to biologically relevant concentrations of immune elicitors, 2) capable of rapid, low cost, and visual assessment of immune activation, and 3) robust to low numbers of biological replicates and background immune elicitation by *Agrobacterium.* Therefore, we utilized the FBP system to develop a reporter of immune activation by heterologously expressed cell surface receptors.

## Materials and Methods

### Plant Materials and Growth Conditions

*N. benthamiana* plants were transplanted one week after sowing and grown at 20°C under 12-hour light and dark cycles. The seedlings were grown under humidity domes for four weeks, after which the domes were removed, and the plants were grown an additional week before infiltrations. Fully expanded, mature leaves of six-week-old plants were used for all transient expression experiments.

### Transcriptomic and qRT-PCR Analysis

For RNAseq analysis, an *N. benthamiana* stable transgenic line expressing *Phaseolus vulgaris* INR (INR-Pv 1-5) (Steinbrenner et al., 2020) was syringe infiltrated with Agrobacterium GV3101 (pMP90) at OD = 0.45 expressing empty vector (EV) pEarleyGate103 (Earley et al., 2006). 24 hpi, *Agrobacterium*-treated leaves were further infiltrated with H_2_O or 1 μM In11 peptide and harvested after an additional 6 hpi. Total RNA was extracted using Nucleospin Plant RNA kit (#740949.250 Macherey-Nagel). RNA was used to generate Lexogen Quantseq 3’ RNA seq libraries at Cornell University Institute of Biotechnology Genomics Facility. 3’ reads were mapped to *N. benthamiana* genome v1.0.1 (Sol Genomics Network) using HISAT2 (Kim et al., 2019) with options min-intronlen 60--max-intronlen 6000, counts by gene were analyzed using HTSeq-Count (Anders et al., 2015) with options -m intersection-nonempty --nonunique all, and differential expression was analyzed by DESeq2 (Love et al., 2014).

For qRT-PCR analysis, *N. benthamiana* plants were syringe infiltrated with *Agrobacterium* (OD_600_ .45) carrying either *p35s::PvINR* or pGreenII empty vector. 24 hours after infiltration, tissue was treated with either water or In11 and harvested after 6 hours. Total RNA was extracted using Trizol reagent (#15596018 Thermo Fisher Scientific, USA). cDNA libraries were generated using SuperScript IV Kit (#18090050 Thermo Fisher Scientific, USA). qRT-PCR reactions were conducted using Applied Biosystems PowerUp SYBR Green Master Mix (#A25742 Thermo Fisher Scientific, USA) and gene specific primer pairs (Supplementary Table S2). Changes in gene expression between water and In11 treatments were calculated using the ΔΔCq method, using ΔCq values normalized against *N. benthamiana EF1α* (D. Liu et al., 2012). Student’s t-tests were performed between comparisons of *35s::PvINR* and EV treated tissue using the ggplot2 package in R (v4.1.2).

### Generation of Reporter Constructs

Promoter regions of candidate marker genes were amplified from genomic DNA of *N. benthamiana* using primers designed against Niben v1.0.1 (Bombarely et al., 2012) with appended overhangs encoding either BsaI or BpiI restriction enzyme recognition sites (Supplementary Table S2). These primers amplified from the start codon to approximately 1.5 kb upstream. Promoter regions were then cloned into the Promoter + 5’ untranslated region (UTR) acceptor backbone obtained from the Golden Gate MoClo Plant Toolkit (Engler et al., 2014). Double promoter constructs were constructed by reamplifying the promoter region of interest with unique overhangs and cloning into the Level −1 universal acceptor backbones using the BsaI-HFv2 restriction enzyme (#R3733L New England Biolabs, USA). These parts were then assembled into the same Promoter + 5’ UTR acceptor backbone using the BpiI restriction enzyme (#ER1012 Thermo Fisher Scientific, USA).

Reporter constructs were generated by first modifying the P307-FBP_6 constitutive autoluminescence construct previously described (Khakhar et al., 2020). P307-FBP_6 was a gift from Daniel Voytas (Addgene plasmid # 139697; http://n2t.net/addgene:139697; RRID:Addgene_139697). To simplify the process of cloning new reporter constructs with promoter regions of interest, the CaMV35s promoter originally used to drive *LUZ* was replaced with an insert encoding a blue-white selectable marker flanked by BsaI recognition sites supplying Promoter + 5’ UTR MoClo overhangs (Supplementary Fig. S2A). This allows for simple, one-step assembly reactions. The promoter regions of interest were then cloned into this acceptor plasmid using the BsaI-HFv2 restriction enzyme. A template primer pair for amplification and cloning putative promoter regions directly into the pFBP_promoter_acceptor construct has been included (Supplementary Table S2).

Reporter constructs were transformed by electroporation into *Agrobacterium tumefaciens* GV3101 (pMP90). All sequences were verified by Sanger sequencing.

### *Agrobacterium-Mediated* Transient Transformation and PAMP treatment

*Agrobacterium* strains carrying the constructs of interest were cultured in LB media containing kanamycin (50μg/mL), gentamicin (50μg/mL), rifampin (50μg/mL), and tetracycline (10ug/mL) for 24h. 3 mLs of culture were then pelleted and resuspended in infiltration media containing 10mM MES (pH 5.6), 10 mM MgCl2, and 150 μM acetosyringone. For coinfiltrations, separate strains harboring reporter and receptor constructs were combined at a final individual OD_600_ = 0.3 for a final cumulative OD_600_ =0.6. After 3h of incubation at room temperature (RT), the cell mixture was infiltrated into fully expanded leaves of 6-week-old *N. benthamiana* plants using a needless syringe.

To assess induction of luminescence, transformed regions were infiltrated with peptide 48 hours after *Agrobacterium* infiltration. Six hours after treatment, leaves were removed at the petiole and luminescence was immediately imaged using the Azure Imaging System with 8 seconds of exposure. Peptides were obtained from Genscript and diluted to specified concentrations in sterile autoclaved water.

### Quantification and Statistical Analysis

Mean gray values of manually defined regions of interest were measured in ImageJ 1.53k. Average signal intensity (ASI) was determined by subtracting the average mean gray value of the untransformed background from the mean gray values of the regions of interest. Negative ASI indicates lower mean gray value than background. One-way ANOVAs and post-hoc Tukey’s t-tests were conducted using the agricolae (v1.3-5) package in R (v4.1.2) and summarized as compact letter displays. Differing letters represent statistically significant differences (p<.05) among pairwise comparisons. Figure editing and layouts were completed in Inkscape.

### Phylogenetic Analysis

Using the annotated coding sequence of *Niben101Scf06684g03003.1*, a BLASTN search was conducted against the *Vigna unguiculata* (v1.2), *Phaseolus vulgaris* (v2.1), and *Arabidopsis thaliana* (TAIR10) genomes (predicted cDNA sequences). *Arabidopsis thaliana* was included to identify potential characterized homologs, and the two legume species were included as representative legume species that natively encode INR. After aligning the top 70 hits, a maximum likelihood phylogenetic tree was generated using FastTree. A subset of this tree was then selected and realigned as translated amino acid sequences using MAFFT (Katoh et al., 2019; Kuraku et al., 2013). A maximum likelihood tree was subsequently generated on the CIPRES web portal using RAXML-HPC2 on XSEDE (v8.2.12) (Miller et al., 2010; Stamatakis, 2014) with the automatic protein model assignment algorithm using maximum likelihood criterion and 100 bootstrap replicates. The resulting phylogeny was rooted and visualized using MEGA11 and edited in Inkscape.

## Results

Differentially Expressed Genes in Response to Agrobacterium and PTI activation Heterologous expression of PRRs in *N. benthamiana* allows for activation of PTI in response to cognate PAMPs, but transient expression requires introduction of *Agrobacterium,* a potentially independent source of PAMPs and activator of PTI responses. To characterize the transcriptional landscape of PTI induced by both *Agrobacterium* and individual PAMP treatment, we conducted transcriptomic analysis in plants stably expressing the *P. vulgaris* Inceptin Receptor (PvINR), an LRR-RLP which recognizes the peptide elicitor inceptin11 (In11) (Steinbrenner et al., 2020). Plants were infiltrated with *Agrobacterium* to mimic conditions during *Agrobacterium-mediated* transient transformation. After 24 hours, the *Agrobacterium-infiltrated* leaves were subsequently treated with water or In11 to induce immune signaling (Supplementary Fig. S1, “AH” or “AI”). Additionally, leaves previously mock infiltrated were infiltrated with water to account for effects of wounding during infiltration (Supplementary Fig. S1, “H”). Tissue was collected after 6 hours, and RNA sequencing was subsequently conducted to identify differentially expressed genes (DEGs) under each pair of conditions.

Compared to leaf tissue not previously infiltrated, infiltration with *Agrobacterium* affected expression of hundreds of genes (Fig. 1A, comparisons “AH vs H” and “AI vs H”, Supplementary Table S1). A total of 1425 upregulated and 938 downregulated genes were significantly altered by *Agrobacterium* infiltration. The majority of DEGs were observed in both In11 and water treated tissue.

**Figure 1.**
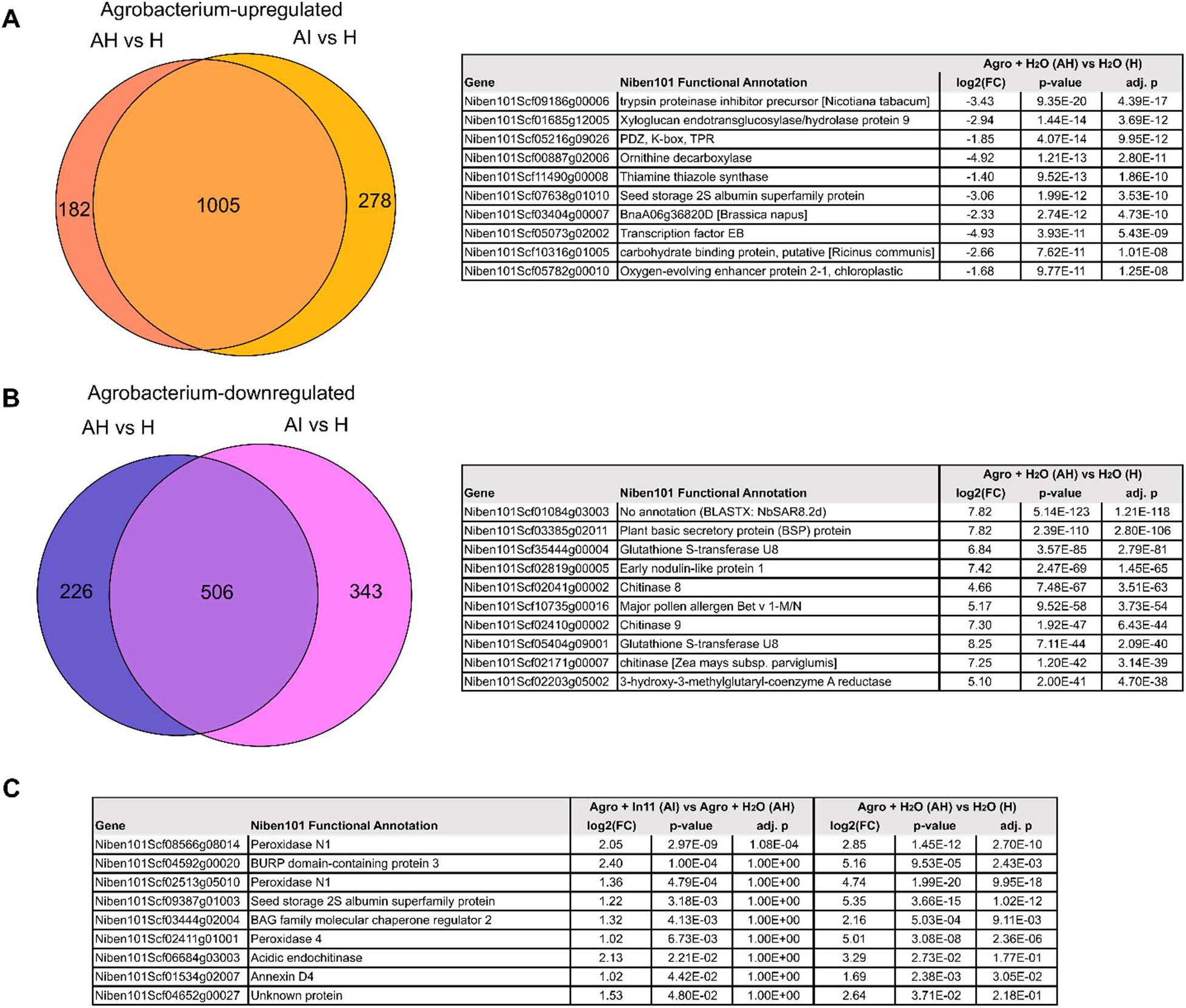
Agrobacterium and In11-induced changes in *N. benthamiana* gene expression. A, Venn diagram displaying number of significantly differentially expressed genes (DEGs) upregulated by Agrobacterium relative to mock-treated tissue. B, Agrobacterium-downregulated genes. Treatments are labeled as follows: AH, Agrobacterium + H_2_O, AI, Agrobacterium + In11, H, Mock infiltrated leaf tissue. See Fig. S1 for treatment details. Top ten genes in both categories with largest log2(fold-change) (FC) are displayed at right. P-value indicates statistical significance with standard Wald test. Adj. P indicates significance after correction for multiple comparisons (Benjamani-Hochberg, BH). C, Candidate genes induced by In11 in the presence of Agrobacterium. While only one DEG was observed after BH correction, 9 genes were induced by In11 uncorrected for multiple comparisons (Fig. S1).

To identify useful markers of PTI activation in the context of *Agrobacterium,* we next compared gene expression in *Agrobacterium-infiltrated* leaf tissue in the presence or absence of In11 peptide (AI vs AH). Only one gene was significantly differentially expressed (Supplementary Table S1, column “adj. p”). Since In11 treatment previously activated measurable early immune phenotypes (Steinbrenner et al., 2020), namely induced ROS and ethylene production, in identical experimental conditions, we reasoned that transcriptional changes at this timepoint may occur below the threshold for statistical significance. We therefore performed a separate analysis filtering for genes with p<0.05 differential expression by standard Wald test but without correction for multiple comparisons (Supplementary Fig. S1B, Supplementary Table S1, column “p-value”). With this relaxed threshold91 genes were characterized as upregulated by the addition of In11 (Supplementary Fig. S1). Interestingly, In11-upregulated genes overlapped with both *Agrobacterium-upregulated* (Supplementary Fig. S1B) and downregulated genes (Supplementary Fig. S1C), suggesting complex regulation of specific *N. benthamiana* PTI outputs.

We further filtered candidate PTI marker genes based on broad responsiveness to both *Agrobacterium* and In11. Nine genes showed higher Agrobacterium or In11 induced expression in all three comparisons (Fig. 1C, Supplementary Fig. S1B), suggesting a large dynamic range of gene expression able to be activated by both *Agrobacterium* PAMPs and by the addition of the separate individual PAMP, In11. To determine more confidently which of these genes are induced by In11 treatment, we conducted qRT-PCR analysis probing differences in expression of each of the candidate marker genes six hours after water and In11 treatment. We found that four genes showed significant induction after treatment with In11 in tissue transiently expressing *PvINR,* but not in tissue infiltrated with an empty vector strain: *Niben101Scf08566g08014, Niben101Scf04592g00020, Niben101Scf06684g03003, Niben101Scf04652g00027* (Fig. 2). We conclude that these four genes serve as markers of INR-mediated responses to In11 in *Nicotiana benthamiana* after transient PRR expression. In summary, while the transcriptional effects of an additional PAMP, In11, 24 hours after Agrobacterium infiltration are subtle, candidate genes were observed with Agrobacterium and PAMP-inducible behavior consistent with broadly responsive marker genes.

**Figure 2.**
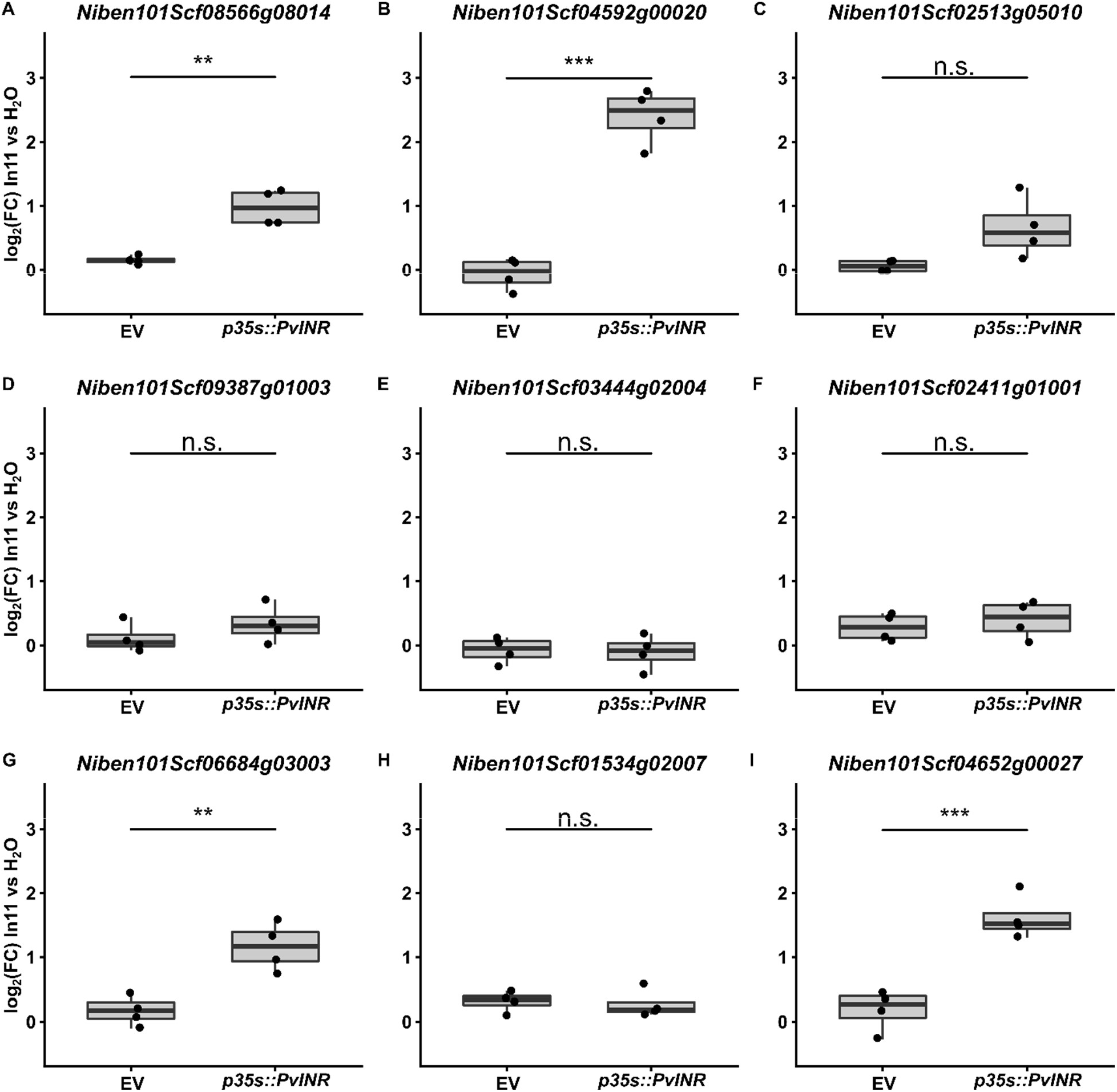
RT-qPCR validation of candidate marker genes. Boxplots indicate the mean log2(fold change) between water and In11 treated tissue (log_2_(FC) In11 vs H_2_O) of gene expression from *N. benthamiana* plants expressing either p*35s::PvINR* or an empty vector (EV) over four biological replicates. Student’s t-tests were conducted to determine significance (n.s.: not significant p>.05, *: p<0.05, **: p<0.01, ***: p<0.001).

### An FBP Luminescence Reporter to Quantify Innate Immune Activation by PTI

To test whether the promoter regions of these genes could function in In11 -inducible reporters, we generated promoter fusion constructs with promoter regions of the endogenous *N. benthamiana* marker genes driving expression of the fungal luciferase (LUZ). Original FBP constructs contain five genes of the pathway for both LUZ and substrate biosynthesis enzymes (Khakhar et al., 2020). We first generated an adaptable acceptor construct allowing MoClo-compatible cloning of promoters to drive LUZ expression (Supplementary Fig. 2A). This vector, *pFBP_promoter_acceptor,* is available on Addgene (confirmation pending). Using this construct, we replaced the original constitutive CaMV35S promoter originally driving LUZ with promoter regions of genes of Fig. 2. Of the two constructs we successfully generated, only a fusion driven by the promoter region of *Niben101Scf06684g03003* showed induction of luminescence upon In11 treatment in a PvINR-dependent manner (Fig. 3A, Supplementary Fig. S3). *Niben101Scf06684g03003* is a homolog of the *A. thaliana* Class III lysozyme *LYS1* (Supplementary Fig. S4) (X. Liu et al., 2014). We therefore termed this construct *pFBP_NbLYS1::LUZ* (hereafter *pLYS1::LUZ).* Importantly, infiltration damage during treatment with H_2_O does not result in high background luminescence and is instead comparable with tissue not infiltrated with the reporter (Supplementary Fig. S5A), making luminescence upon induction with In11 easily detectable.

**Figure 3.**
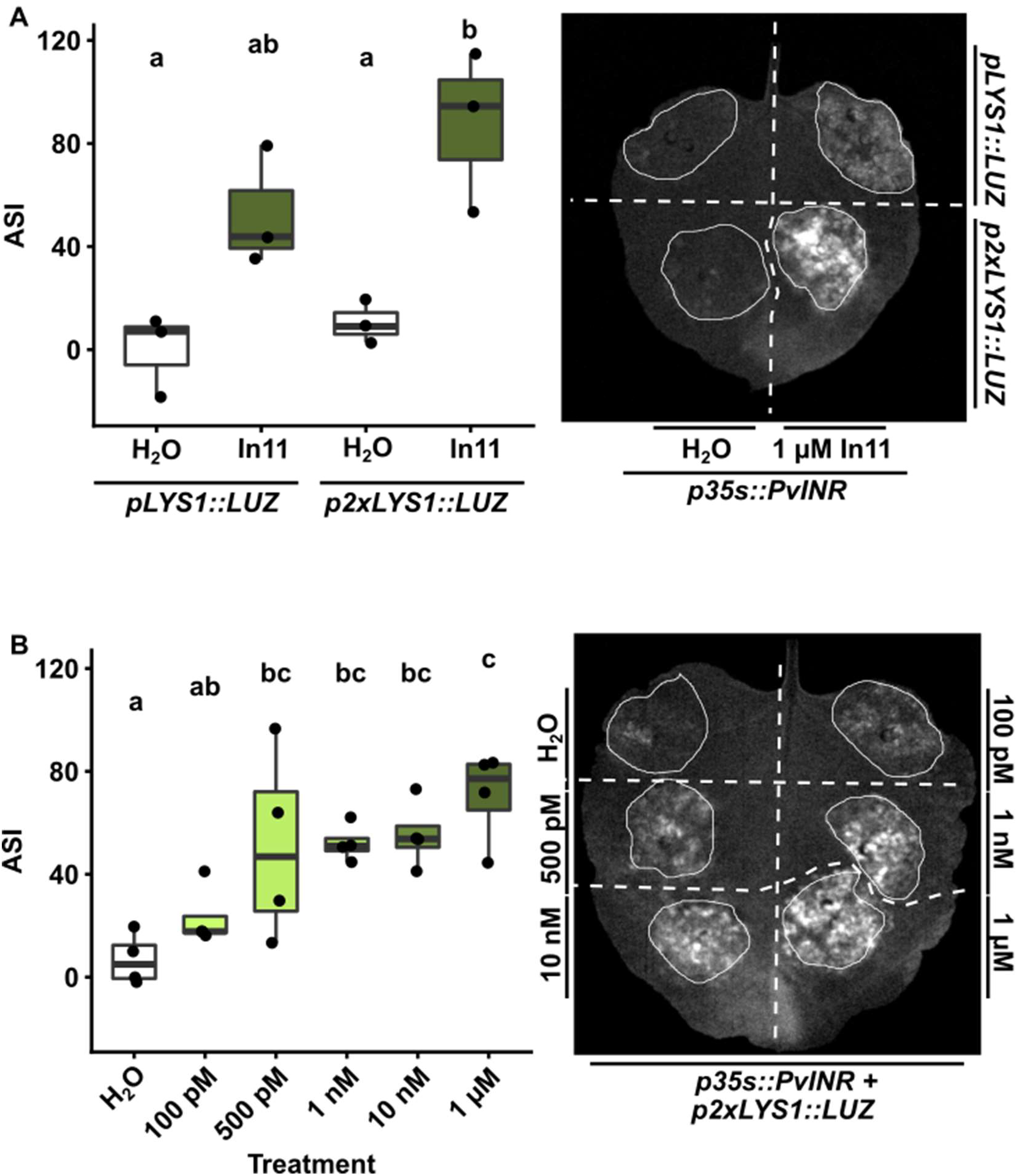
A *Nicotiana benthamiana LYS1* homolog serves as a marker of inceptin response. A, Leaves were coinfiltrated with *p35s::PvINR* and *pLYS1::LUZ* across the proximal portion of the leaf, and *p35s::PvINR* and *p2xLYS1::LUZ* across the distal portion of the leaf. 48 hours after infiltration, one half of the leaf was infiltrated with sterile water, and the other half was infiltrated with 1 μM In11. Images were obtained 6 hours after peptide treatment, and ASI was quantified in ImageJ. Left, boxplots show the average ASI of three independent biological replicates. Letters represent significantly different means (One-way ANOVA and post-hoc Tukey’s HSD tests, p<.05). Right, a representative leaf image of one biological replicate is depicted. B, Leaves were co-infiltrated with *p35s::PvINR* and *p2xLYS1::LUZ* in six distinct regions of the leaf. 48 hours after infiltration each zone was infiltrated with sterile water or a series of In11 concentrations. Imaging and quantification were conducted as in A.

Luminescence induced by the *pLYS1::LUZ* construct was markedly lower than a construct using the CaMV35S promoter to drive LUZ expression (Supplementary Fig. S4B). However, duplication of promoters has been shown to effectively increase strength of expression (Kay et al., 1987). To enhance the strength of reporter expression and observable luminescence, we also constructed *pFBP_2xNbLYS1::LUZ* (hereafter *p2xLYS1::LUZ),* a double promoter region construct where two copies of the *NbLYS1* promoter were arranged adjacently and used to drive expression of luciferase treatment (Supplementary Fig. S5B). When coexpressed alongside *PvINR,* the *pLYS1::LUZ* single copy construct did not show a statistically significant difference in luminescence between water and In11 treatment, while the *p2xLYS1::LUZ* double copy construct did show a significant difference between the water and In11 treatments (Fig. 3A). We therefore elected to proceed using the *p2xLYS1::LUZ* FBP construct as a reporter for all subsequent experiments.

Different assays for immune receptor function show varying degrees of sensitivity to low elicitor concentrations (Mott et al., 2018). To determine the sensitivity of the FBP reporter assay to In11 treatment, we coexpressed the *p2xLYS1::LUZ* reporter and *p35s::PvINR* and conducted a dose-response experiment using increasing concentrations of In11. We observed statistically significant differences in luminescence between water and In11 treatments above 500 pM (Fig. 3B). This falls within the range of reported In11 concentrations that are present in the oral secretions of caterpillars during herbivory (Schmelz et al., 2006). As a result, the *p2xLYS1::LUZ* is a robust reporter of immune activation by biologically relevant elicitor concentrations.

Besides INR, other heterologously expressed PRRs are capable of conferring PTI immune signaling in *N. benthamiana* (Albert et al., 2015; Steinbrenner et al., 2020; L. Zhang et al., 2021). Furthermore, flg22 treatment induces expression of the *A. thaliana LYS1* homolog (X. Liu et al., 2014). To test whether the *p2xLYS1::LUZ* construct serves as a reporter of PRR activity more broadly, we also tested reporter inducibility by two cell surface PRRs from *A. thaliana:* EFR and RLP23. We observed background induction of luminescence in leaf tissue expressing EFR when treated with elf18 (Fig. 4C), potentially due to background induction of EFR by *Agrobacterium.* elf18 nonetheless robustly induces luminescence relative to mock treatment. We also observed induction of luminescence in leaf tissue expressing RLP23 when treated with nlp20 (Fig. 4D). Importantly, induction of luminescence is only observed in regions of interest where the cognate receptor-elicitor pair is present. Together, these data support the utility of the *p2xLYS1::LUZ* construct as a robust reporter of specific PRR-elicitor interactions.

**Figure 4.**
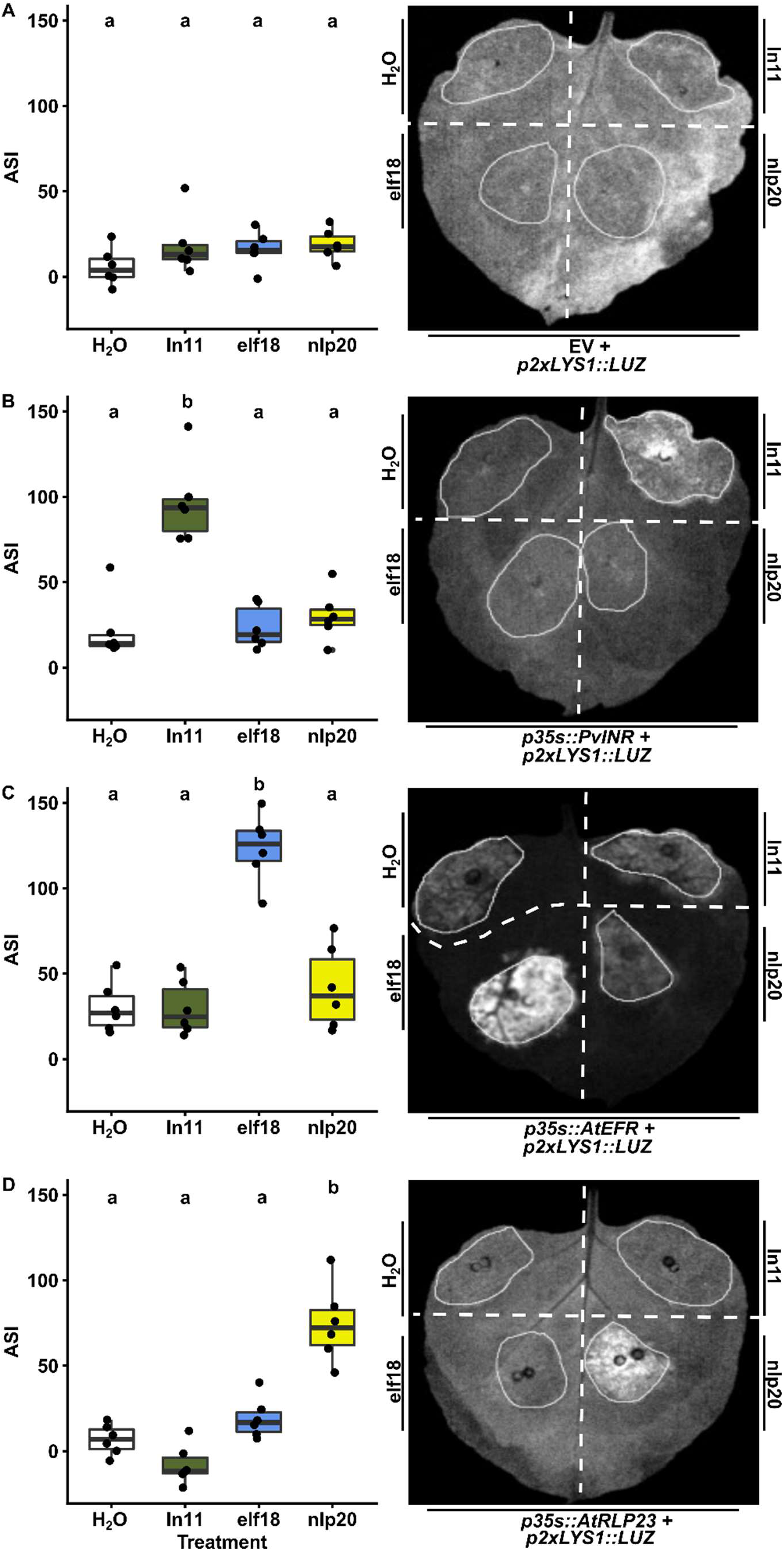
The *p2xLYS1::LUZ* construct acts as a generic reporter for plant pattern recognition receptors. Leaves of *N. benthamiana* plants were coinfiltrated with the *p2xLYS1::LUZ* reporter construct and either A) empty vector, B) *35s::PvINR,* C) *35s::AtEFR,* or D) *35s::AtRLP23* in four distinct regions. 48 hours after infiltration, each region was infiltrated with either sterile water, 1 μM In11, 1 μM elf18, or 1 μM nlp20 peptide. Images were obtained 6 hours after peptide treatment, and ASI was quantified in ImageJ. Left, boxplots show the average ASI of six independent biological replicates. Letters represent significantly different means (One-way ANOVA and post-hoc Tukey’s HSD tests, p<.05). Right, a representative leaf image of one biological replicate is depicted.

### SOBIR1 is necessary for INR-mediated Induction of Bioluminescence by Inceptin

Characterized LRR-RLPs are known to require the adaptor receptor-kinase SUPPRESSOR OF BIR1-1 (SOBIR1) to initiate downstream signaling (Liebrand et al., 2013; Albert et al., 2015). Although SOBIR1 has been shown to associate with INR in *Nicotiana benthamiana,* it is not yet known if SOBIR1 is necessary for immune signaling by PvINR (Steinbrenner et al., 2020). To determine whether PvINR requires SOBIR1 and whether the *p2xLYS1::LUZ* reflects downstream immune signaling pathways, we conducted reporter assays in *N. benthamiana sobir1* knockout plants, which previously showed compromised function of the tomato LRR-RLP Cf4 (Huang et al., 2021). Induction of luminescence by In11 treatment is absent in *sobir1* mutant plants and restored when either *A. thaliana SOBIR1* or *P. vulgaris SOBIR1* are coexpressed with *PvINR.* (Fig. 5). Thus, reporter activation is subject to similar requirements for LRR-RLP function as well-characterized PTI responses. This suggests that the *p2xLYS1::LUZ* construct serves as a useful tool not only for studying receptor-elicitor interactions but also downstream interactions important for immune activation and signaling.

**Figure 5.**
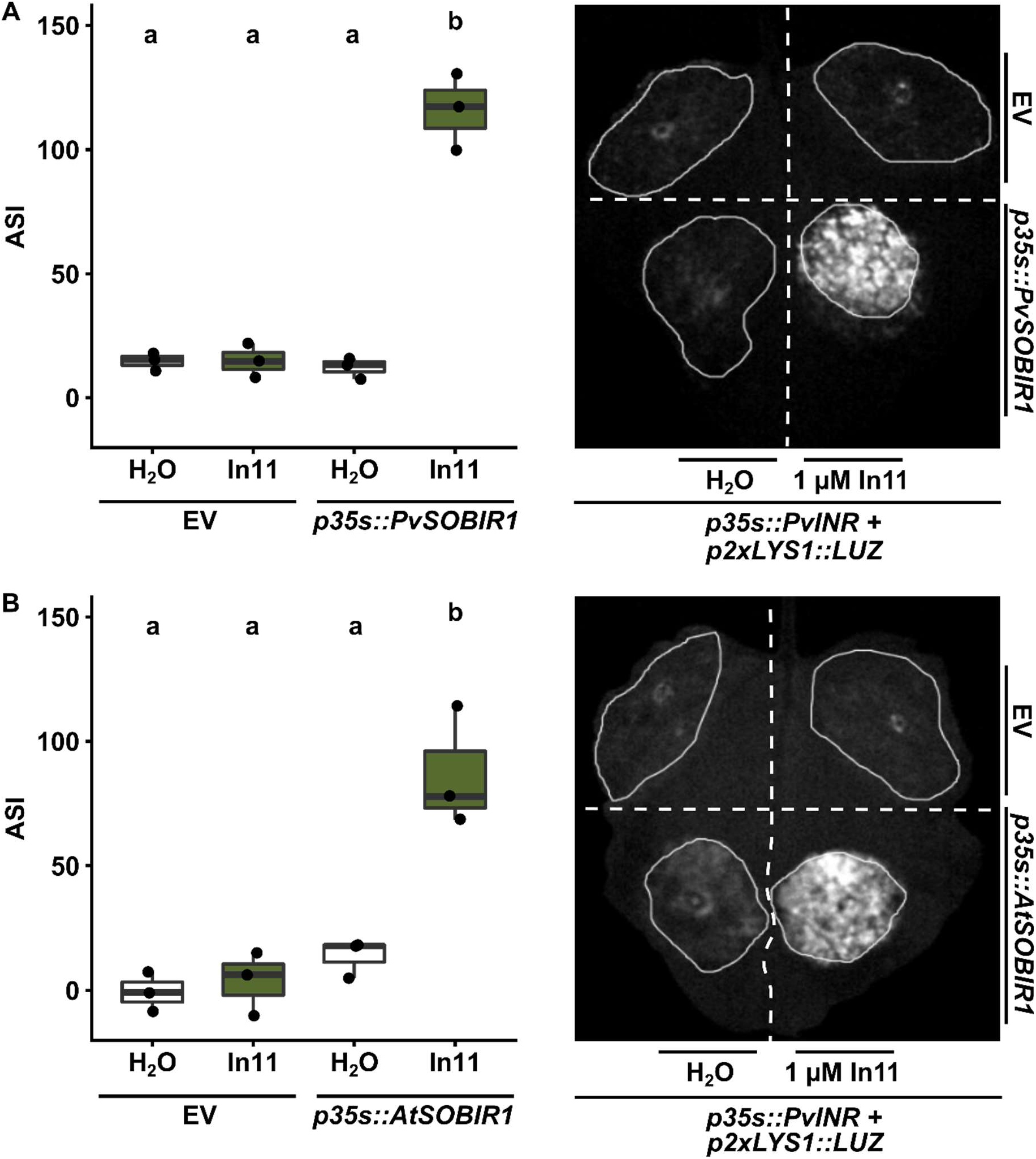
SOBIR1 is necessary for activation of luminescence by PvINR. Leaves of *Nicotiana benthamiana sobir1* knockout plants were coinfiltrated with the *pFBP_2xLYS1::LUZ* reporter construct, *p35s::PvINR* and either: empty vector (EV); A) *p35s::PvSOBIR1; or B) p35s::AtSOBIR1*, repeated for three biological replicates. 48 hours after infiltration, each region of interest was infiltrated with either sterile water or 1 μM In11. Images were obtained 6 hours after peptide treatment, and ASI was quantified in ImageJ. Left, boxplots show the average ASI of three independent biological replicates. Letters represent significantly different means (One-way ANOVA and post-hoc Tukey’s HSD tests, p<.05). Right, a representative leaf image of one biological replicate is depicted.

## Discussion

We describe here a genetically encoded reporter responsive to heterologously expressed PRRs in *N. benthamiana.* The *p2xLYS1::LUZ* reporter demonstrates robust PAMP sensitivity and does not require addition of exogenous enzyme substrate. Therefore, this reporter assay may be a useful tool for assessing immune activation by a number of diverse PRRs, including both receptor-like kinases and receptor-like proteins.

To develop this reporter, we first characterized the transcriptional modifications that occur in response to both *Agrobacterium* and elicitor perception by a heterologously expressed LRR-RLP. LRR-RLPs warrant further structural and functional characterization, as they constitute a key class of PRRs involved in activating plant innate immune responses (Jamieson et al., 2018; Albert et al., 2020; Steinbrenner, 2020). LRR-RLPs also include the first known receptor-ligand pair involved in defense against a chewing herbivore (Steinbrenner et al., 2020). However, the specific molecular interactions required for immune signaling by LRR-RLPs in plants remain only partially understood, in part because no solved crystal structures of LRR-RLPs have been reported. Although characterized LRR-RLPs require SUPPRESSOR OF BIR1-1 (SOBIR1) and SOMATIC EMBRYOGENESIS RECEPTOR KINASEs (SERKs) to activate immune signaling, the mechanisms underlying ligand binding and coreceptor association are unclear (van der Burgh et al., 2019). As a result, we tailored our reporter system toward heterologously expressed LRR-RLPs to aid in gathering deeper insights into this important and incompletely understood group of plant cell surface immune receptors. However, we also observed induction of luminescence in response to the bacterial elicitor elf18 in tissue expressing *A. thaliana* EFR, a receptor-like kinase (RLK) (Zipfel et al., 2006). As a result, this reporter could be useful for studying RLK signaling. Additionally, recent studies describing the overlap between PTI and ETI signaling suggest common signaling components (Ngou et al., 2021; Pruitt et al., 2021). As a result, there is a possibility this reporter could serve to study intracellular immune receptors and may be particularly useful when these receptors do not produce hypersensitive responses.

Unsurprisingly, our transcriptomic analysis revealed that *Agrobacterium* treatment alone resulted in large changes in gene expression. This demonstrates that *Agrobacterium* strongly induces innate immunity in *N. benthamiana,* likely through recognition of *Agrobacterium* PAMPs, resulting in large-scale transcriptional changes. Therefore, it is important to consider the role of *Agrobacterium* PAMPs in activating immunity. Interestingly, many genes that showed upregulation in response to In11 treatment were genes that were downregulated by *Agrobacterium* (Fig S1B-C). While likely due to timescales of *Agrobacterium* inoculation (24 hpi) versus In11 treatment (6 hpi), it is also possible that perception of Agrobacterium PAMPs by *N. benthamiana* is antagonized by simultaneous activation of immunity by the herbivore-associated In11 elicitor, a potential result of signaling conflict between SA and JA signaling (Li et al., 2019). *Agrobacterium* may activate biotroph-related immunity, whereas INR may activate necrotroph-related immunity through pathways downstream of SOBIR1. Because of the complex nature of these factors, we selected genes that showed upregulation in response to *Agrobacterium* that was further amplified by In11 treatment to identify a generic marker of immune activation by specific elicitor receptor interactions.

Although we identified four marker candidates, only one showed induction of luminescence in response to elicitor treatment (Fig. 3). We were either unable to clone the respective promoter region *(Niben101Scf04592g00020, Niben101Scf04652g00027)* or observed no luminescence in response to In11 treatment compared to water treatment *(Niben101Scf08566g08014)* (Supplementary Fig. S2). Although protocols exist to amplify difficult templates such as AT or GC-rich sequences (Dhatterwal et al., 2017; Sahdev et al., 2007), the complex nature of the *N. benthamiana* genome poses technical challenges in amplifying already evasive promoter regions (Bombarely et al., 2012). Furthermore, it is possible that promoter terminator incompatibility occurred between candidate promoter regions resulting in silencing of *LUZ* (P.-H. Wang et al., 2020). Finally, it is possible that we failed to include the necessary cis-regulatory elements of the promoter region, as we decided on a somewhat arbitrary cutoff of 1.5 kb preceding the start codon of the gene. Trans-regulatory elements may also be necessary to mediate observed changes in gene expression in response to In11 treatment. As a result, improved understanding of plant transcriptional regulatory elements will facilitate efforts to identify and utilize additional highly responsive promoters under a variety of biotic stress conditions as tools to study plant immune responses.

The *pFBP_promoter_acceptor* construct is now publicly available to screen other candidate promoters through simple MoClo ligation of a promoter of interest. However, several considerations should be taken regarding the use of the FBP reporter. Unlike firefly luciferase, which is known to have a short half-life in the presence of luciferin (Van Leeuwen et al., 2000), it is suggested that the stability of fungal luciferase is more suitable for measuring changes over hours, limiting its utility on finer time scales (Khakhar et al., 2020). Induction of luminescence should as a result be viewed as a cumulative representation of reporter activity, rather than instantaneous measure of gene expression. Furthermore, production of fungal luciferin depends on availability of caffeic acid, causing luciferin availability to not be completely uniform across all plant tissues, and sustained periods of fungal luciferase activity to possibly deplete luciferin stores. Although these limitations remain largely negligible for the purpose of assessing specific receptor-elicitor interactions, they should be considered in situations where temporal and spatial aspects are of importance.

This system represents a potentially high-throughput and sensitive reporter for assessing immune activation by heterologously expressed PRRs in *N. benthamiana.* Although other systems retain power by being more sensitive to subtle immune phenotypes and usefulness in characterizing endogenous immune signaling processes in non-model organisms, the sensitivity, robustness, and ease of the FBP reporter system make it useful for understanding cell surface receptor function in *N. benthamiana.* As a result, this reporter represents a potentially valuable addition to the plant immune biology toolkit, especially for large scale studies aimed at illuminating the structure and function of cell surface immune receptors from diverse species.

## Acknowledgements

We thank members of the Steinbrenner, Imaizumi and Nemhauser labs at UW Biology for helpful discussion and comments. This work was supported by NSF grant IOS-2139986 and the UW Mary Gates Endowment for Students.

## Author Contributions

A.D.S. conducted the transcriptomic analysis. A.G.K.G. conducted subsequent experiments. A.D.S. and A.G.K.G wrote the manuscript.

## Materials Availability

Transcriptomic data are available in NCBI SRA (BioProject number pending). The *pFBP_promoter_acceptor* and *pFBP_2xNbLYS1::LUZ* constructs are available at Addgene (confirmation pending).

## Supplementary Figures

**Pg 2. Supplementary Figure S1.** Treatment details and transcriptional changes without correction for multiple comparisons.

**Pg 3. Supplementary Figure S2**. A modified FBP construct for rapid cloning and testing of putative promoter regions.

**Pg 4. Supplemental Figure S3**. The putative promoter region of *Niben101Scf08566g08014* does not serve as a marker of In11 response.

**Pg 5. Supplemental Figure S4**. The In11-upregulated gene *Niben101Scf06684g03003.1* is a homolog of *A. thaliana LYS1.*

**Pg 6. Supplementary Figure S5**. The *LYS1* Promoter Drives *LUZ* weaker than the strong constitutive *CaMV35s* promoter.

**Supplementary Figure S1.**
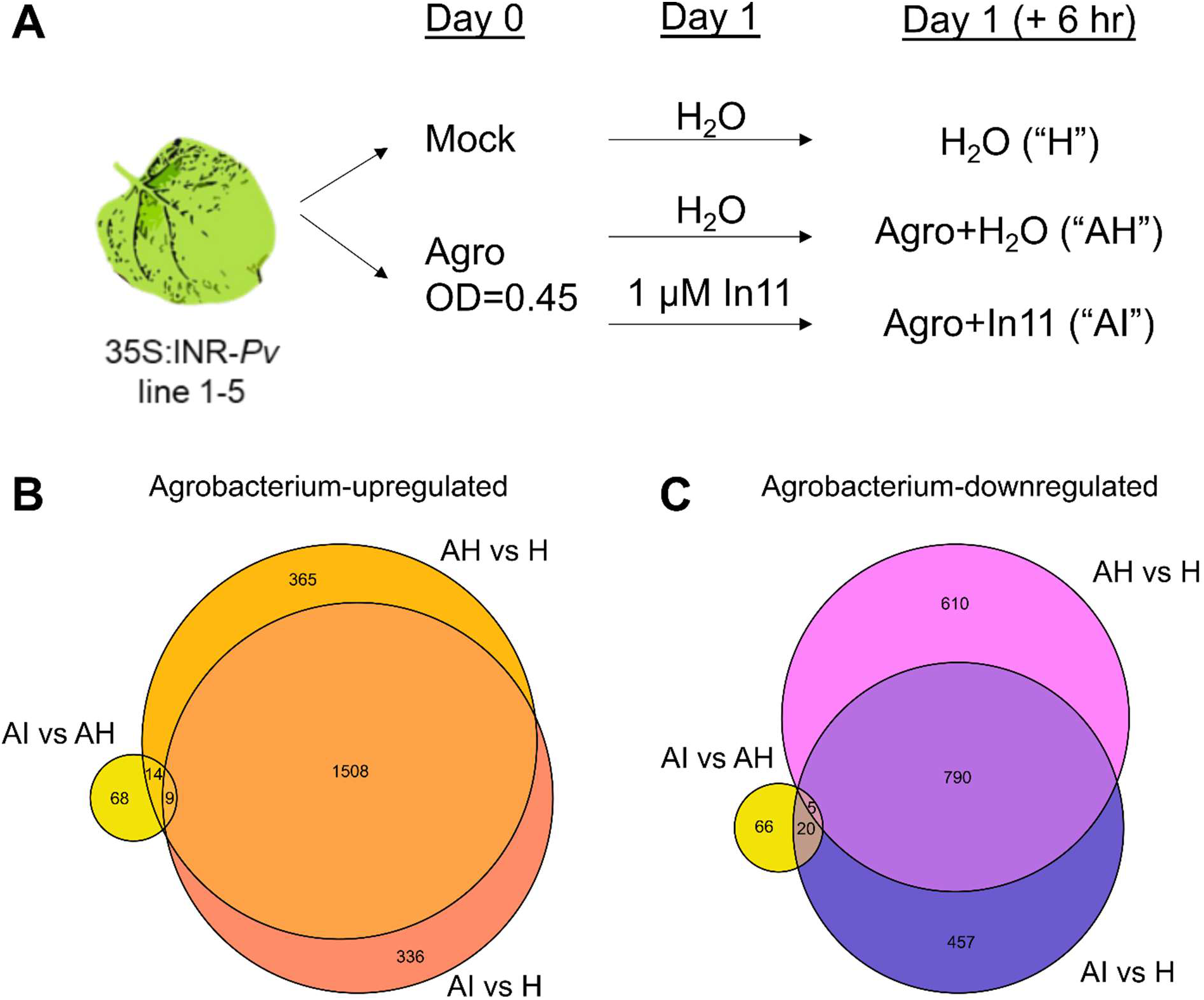
Treatment details and transcriptional changes without correction for multiple comparisons. A, *N. benthamiana* line 1-5 was infiltrated with water (mock) or Agrobacterium and then infiltrated at 24 hpi. Three treatments were collected for RNAseq analysis as indicated. Treatments are labeled as follows: AH, Agrobacterium + H_2_O, AI, Agrobacterium + In11, H, Mock infiltrated leaf tissue. B-C, Venn diagram displaying number of genes upregulated by In11 in the presence of Agrobacterium (AI vs AH, yellow), and genes B) upregulated by Agrobacterium or C) downregulated by Agrobacterium (Wald test, p-value < 0.05, no correction for multiple comparisons by Benjamini-Hochberg).

**Supplementary Figure S2.**
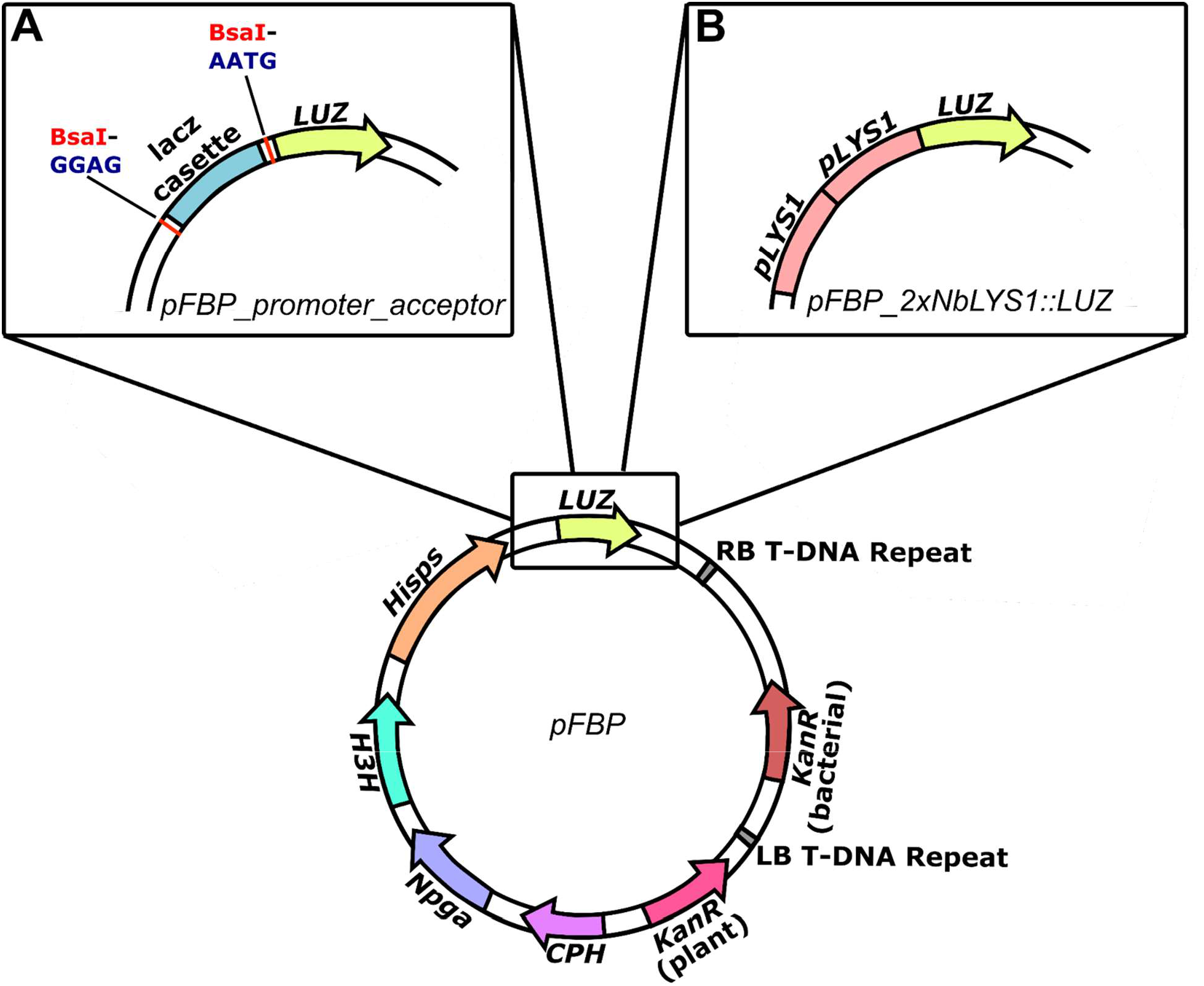
A modified FBP construct for rapid cloning and testing of putative promoter regions. A, The *pFBP_promoter_acceptor* construct contains a *lacz* cassette that allows for bluewhite screening to facilitate screening of correct clones. This cassette is flanked by BsaI recognition sites that exposes GGAG and AATG overhangs upon digestion. These overhangs are compatible with MoClo Promoter +5’ Untranslated Region Level 0 constructs. B, The *pFBP_2xNbLYS1::LUZ* construct places two identical promoter region sequences of a *N. benthamiana* homolog of *A. thaliana* LYS1 immediately adjacent to the coding sequence of *Neonothopanus nambii LUZ.*

**Supplementary Figure S3.**
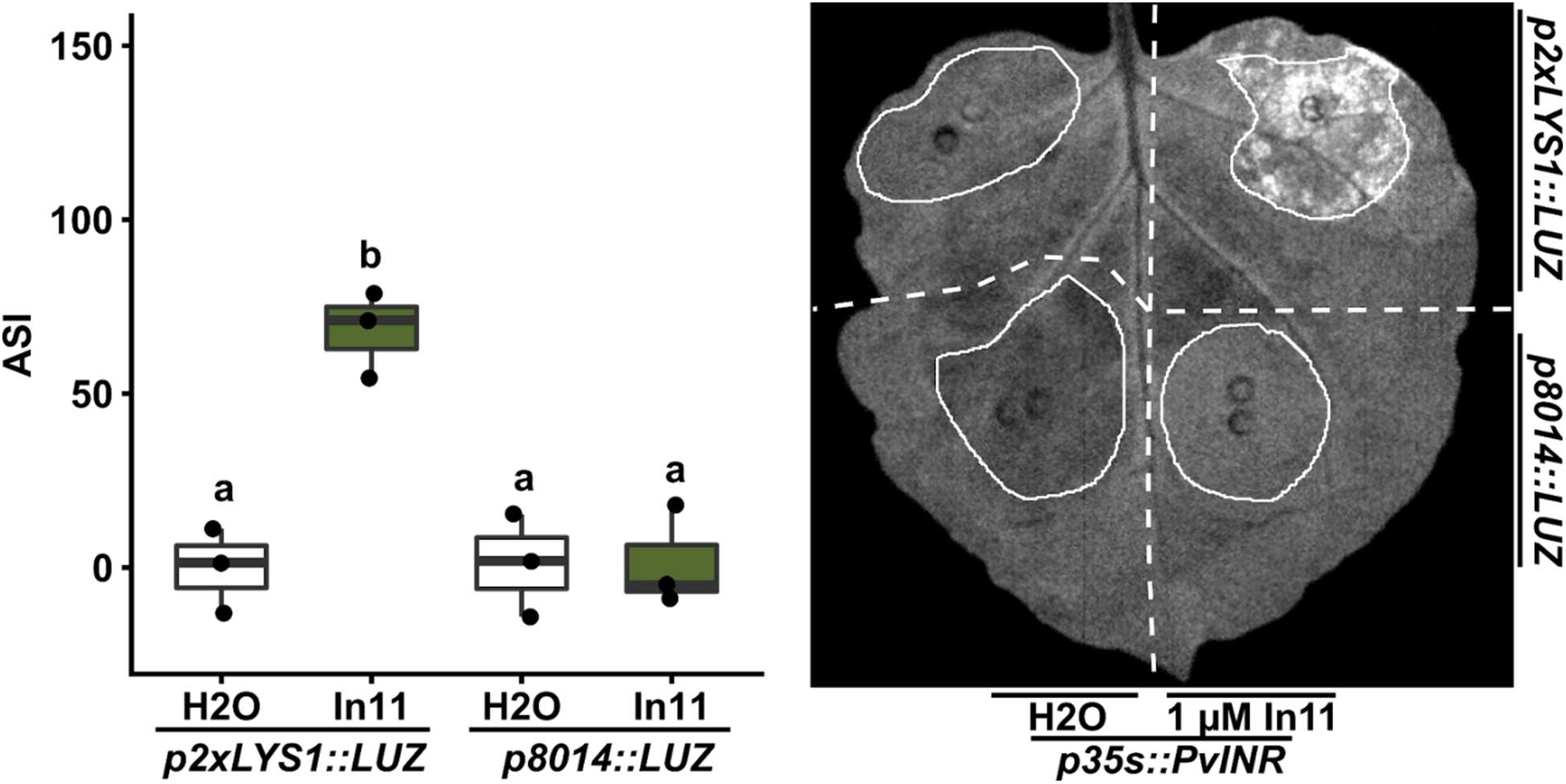
The putative promoter region of *Niben101Scf08566g08014* does not serve as a marker of In11 response. We generated *p8014::LUZ,* a construct using the putative promoter region of *Niben101Scf08566g08014* to drive *LUZ* expression. Leaves were coinfiltrated with *35s::PvINR* and *pLYS1::LUZ* across the proximal portion of the leaf, and *35s::PvINR* and *p8014::LUZ* across the distal portion of the leaf. 48 hours after infiltration, one half of the leaf was infiltrated with sterile water, and the other half was infiltrated with 1 μM In11. Images were obtained 6 hours after peptide treatment, and ASI was quantified in ImageJ. Left, boxplots show the average ASI of three independent biological replicates. Letters represent significantly different means (One-way ANOVA and post-hoc Tukey’s HSD tests, p<.05). Right, a representative leaf image of one biological replicate is depicted.

**Supplementary Figure S4.**
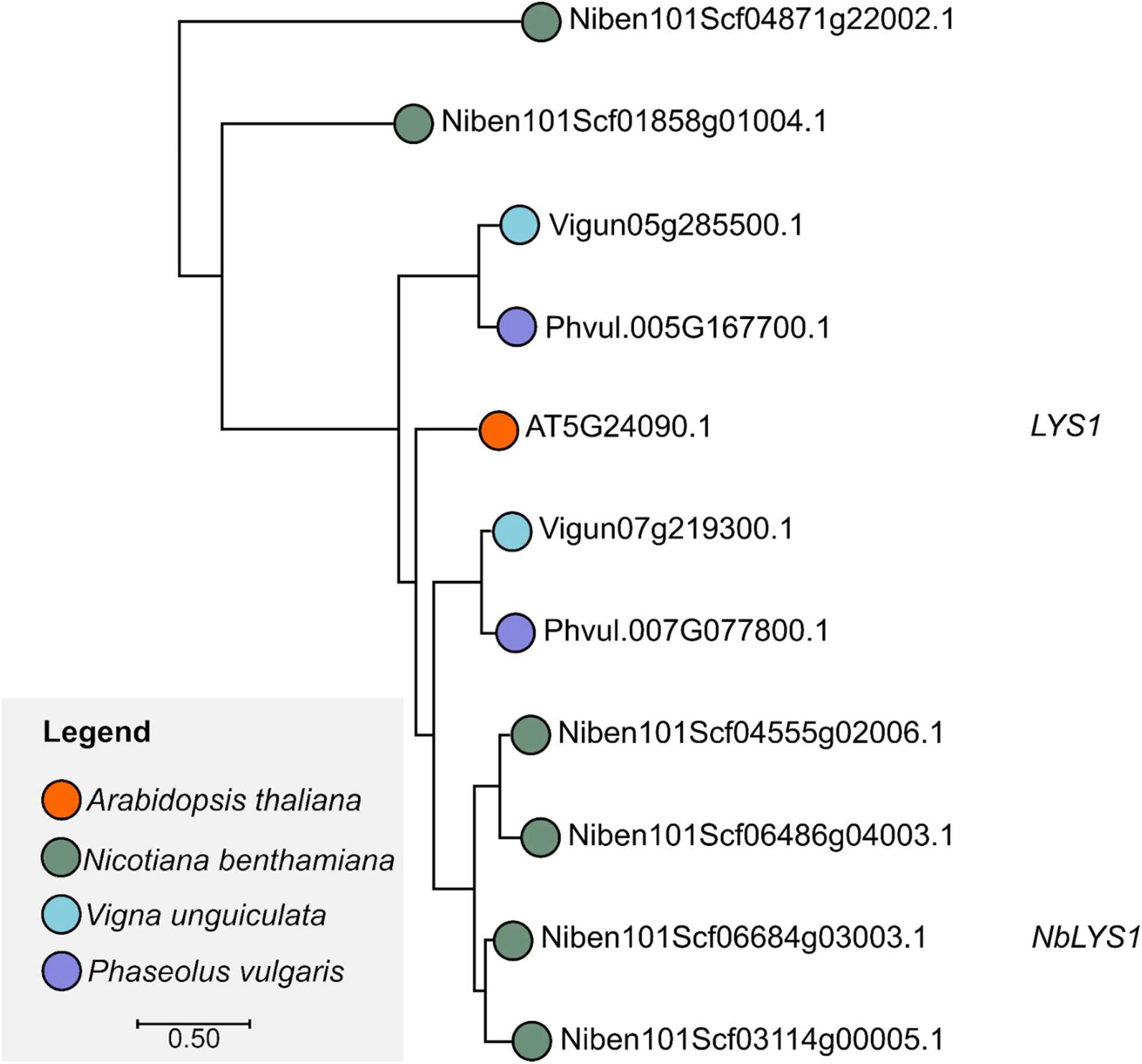
The In11-upregulated gene *Niben101Scf06684g03003.1* is a homolog of *A. thaliana LYS1*. A maximum likelihood tree generated from BLASTN results of the *Niben101Scf06684g03003.1* gene indicates close homology to the characterized *Arabidopsis thaliana LYS1* gene.

**Supplementary Figure S5.**
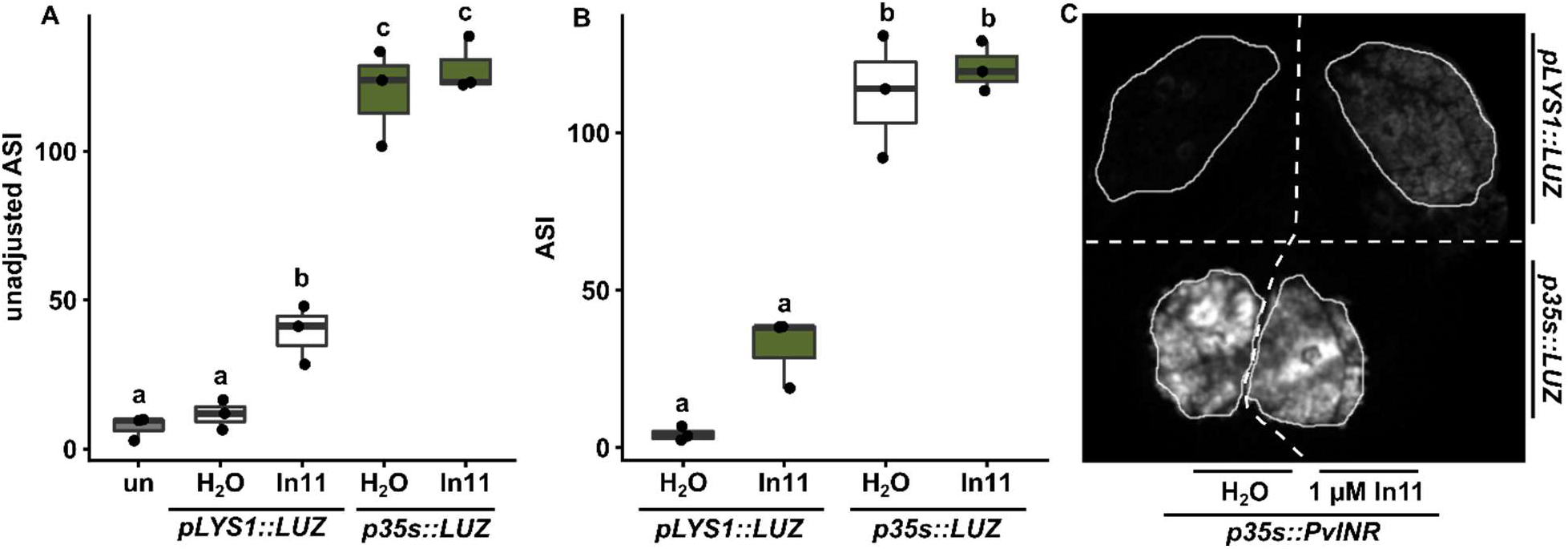
The *LYS1* Promoter Drives *LUZ* weaker than the strong constitutive *CaMV35s* promoter. *N. benthamiana* leaves were infiltrated with *p35s::PvINR* and *pLYS1::LUZ* across the proximal portion of the leaf, and *p35s::PvINR* and *p35s::LUZ* across the distal portion of the leaf. 48 hpi, each half of the leaf was infiltrated with either water of 1μM In11. Images were obtained 6 hours after peptide treatment, and average signal intensity (ASI) was quantified using ImageJ. One-way ANOVA and post-hoc Tukey’s HSD tests were conducted and summarized as a compact letter display. Statistically significant differences (p <.05) among pairwise comparisons are represented by differing letters. A, Unadjusted ASI demonstrates insignificant difference in luminescence produced upon water treatment compared to untransformed and uninfiltrated plant tissue (un). B, Adjusted ASI over three biological replicates reveals relative strengths of expression by the PAMP responsive *NbLYS1* promoter and the strong constitutive *CaMV35s* promoter. C, A representative leaf is shown.

